# Spontaneous emergence of alternating hippocampal theta sequences in a simple 2D adaptation model

**DOI:** 10.1101/2024.06.10.598313

**Authors:** John Widloski, David J. Foster

## Abstract

Spatial sequences encoded by cells in the hippocampal-entorhinal region have been observed to spontaneously alternate across the animal’s midline during navigation in the open field, but it is unknown how this occurs. We show that sinusoidal sampling patterns emerge rapidly and robustly in a simple model of the hippocampus that makes no assumptions about sequence direction. We corroborate our findings using hippocampal data from rats navigating in the open field.

## MAIN

Animals often exhibit alternating left-right search patterns in order to track and localize targets in their environment (Murlis et al., 1992). For example, flying moths that lose an odor plume perform alternating side-to-side “casting” movements in their trajectory in order to regain it (Kennedy & Marsh, 1974), as do rodents that lose an odor trail (Khan et al., 2012). Egyptian fruit bats approaching a target elicit sonar beams that alternate with respect to the body midline at a fixed angular offset, thought to aid in target discrimination and localization (Yovel et al., 2010). These alternating behavioral search patterns are also mirrored at the neural level. In mammals during locomotion, hippocampal place cells, which fire in restricted zones in an environment, activate in sweeps called “theta sequences” that last ∼150 msec and extend forward from the animal’s current location (Foster & Wilson, 2007; Gupta et al., 2012; Wikenheiser & Redish, 2015). At binary decision points, theta sequences sweep cyclically across the two alternatives (Kay et al., 2020), thought to be a virtual form of trial-and-error or active *memory* sampling by which the animal plays out the consequences of actions internally (Redish, 2016). In the open field, grid cells in the medial entorhinal cortex, which are sensitive to both the animal’s location and direction (Hafting et al., 2005; Sargolini et al., 2006) have been shown to encode sweeps during theta oscillations that occur at a fixed angle of approximately 30 degrees but alternate across the body midline (Vollan et al., 2024). Similar to the structure and function of echolocating sonar beams in the bat, alternating theta sweeps in the MEC might be tuned to discern approaching objects in memory space.

How the brain generates patterns of neural activity that are not only sequential but spontaneously alternate has become a recent topic of debate. Most notable are models that marry continuous attractor dynamics, which possess a continuum of stable bump-like states useful for encoding representations of continuous variables like position or head direction with short-term adaptation, which acts to locally depress active neurons within the bump and causing it sweep across the network (Hansel & Sompolinsky, 1997; Hopfield, 2010; Itskov et al., 2011; Azizi et al., 2013; Ecker et al., 2022; Milstein et al., 2023; Chu et al., 2023; Tsodyks et al., 1998; York & van Rossum, 1996; Romani & Tsodyks, 2015). These models naturally account for a wide range of phenomenology related to hippocampal sequences, including the backwards-then-forwards “Nike” checkmark of individual theta sequences on linear tracks (Wang et al., 2020; Chu et al., 2023) and the cyclical sweeping across arms of a T-maze (Romani et al., 2015; Chu et al., 2023). However, it is unclear whether such models, which have only been applied to linearized environments, can spontaneously self-organize to exhibit the oriented, alternating sweeps that are characteristic of theta sequences in 2D. Here, we demonstrate that this simple model can exhibit theta sequences that alternate regularly across the front of the animal, with an angular offset of approximately 30 degrees that is strikingly similar to empirical observations. Importantly, this regularity arises rapidly and robustly in the model with very little fine tuning. Lastly, we confirm our key findings using hippocampal data from rats navigating in the open field.

To model theta sequences in a 2D environment, we simulated stochastic spiking in a sheet of neurons representing pyramidal cells in CA3 (**Figure S1A**), where spiking was modeled as an inhomogeneous sub-Poisson process. The use of a non-deterministic spiking model ensured that any structure in the emergent network dynamics was robust to spiking noise. Neurons were connected with local excitation and global inhibition, a form of recurrence that shapes uniform network excitation into “bumps” of activity that can exist stably anywhere on the sheet. Sweeping dynamics in the bump activity was elicited by including an accumulating (∼1 sec) spike history-based feedback inhibition. This is a form of firing rate adaptation (Hansel & Sompolinsky, 1997; Hopfield, 2010; Itskov et al., 2011; Azizi et al., 2013; Ecker et al., 2022; Milstein et al., 2023; Chu et al., 2023) that mimics the effects of slow calcium-dependent potassium channels. In addition, each neuron received place-like inputs that were unimodal as a function of rat location (**Figure S1A**), with place preferences assumed to uniformly tile the 2D environment. Place inputs were derived from a rat trajectory simulated as a correlated random walk within a 1 m^2^ enclosure (**Figure S1B-C**). Lastly, a fictive, stochastic theta oscillation was imposed on the network dynamics (**Figure S1D**), with theta periods across time sampled from a distribution matched to data (compare **Figure 1F** and **Figure 2F**). Place input was pulsed in the network at the beginning of each theta cycle while the recurrent and adaptive inputs were turned on later in the cycle, mimicking the trading off between sensory-driven and retrieval-driven dynamics thought to occur at different phases of theta (Hasselmo & Stern, 2014). Together, these components yielded sweeps in the bump activity state that emanated outwards from the rat at the beginning of each theta cycle but crucially did not constrain its direction. Adaptation strength was tuned to yield sweeps of approximately 15 cm within each theta cycle so as to match experimental data (**Figure S2A**; compare **Figure 1G** and **Figure 2G**).

**Figure 1:**
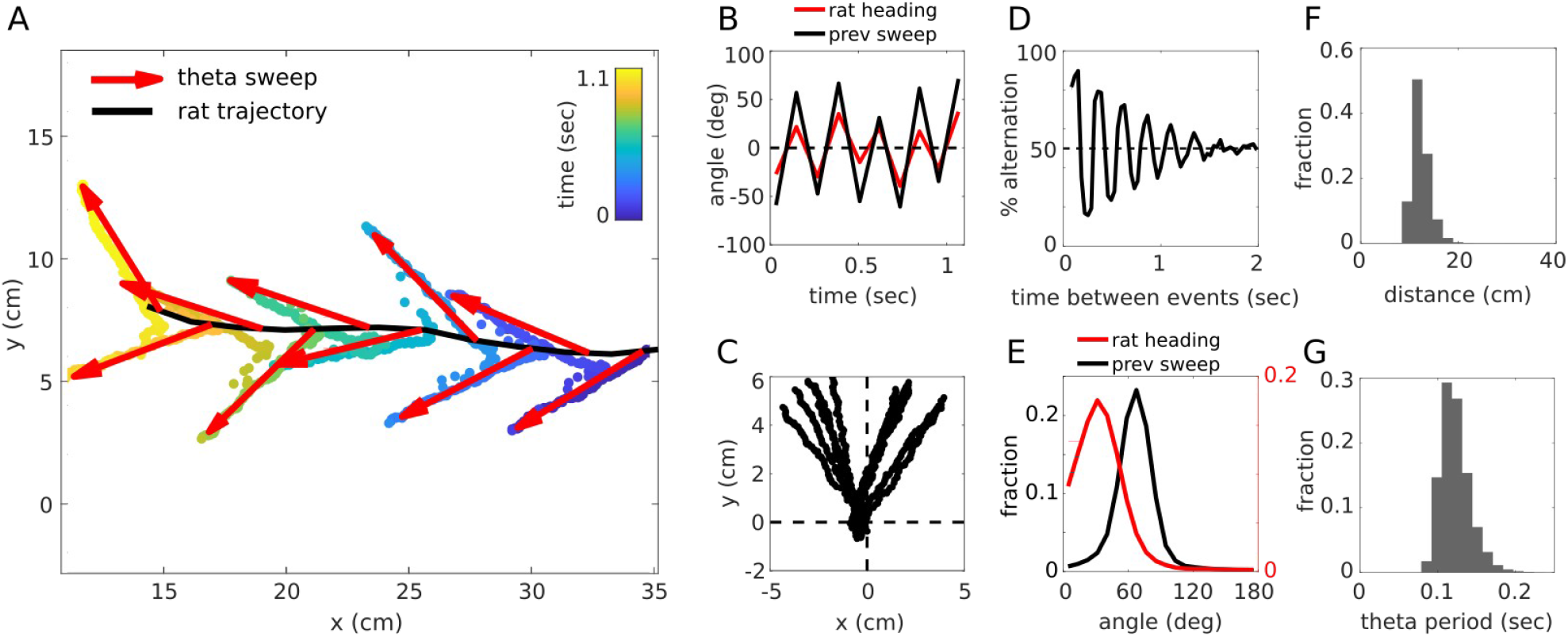
Spontaneous emergence of theta sequence alternation in 2D model. (A) Center-of-mass trajectory of the simulated activity bump depicted across 10 theta cycles, colored according to elapsed time. Red arrow depicts sweep angle, measured as the angle of the line that bisects the rat’s mean location and the location of the sweep at its furthest extent within each theta cycle. Dashed black line is the rat’s trajectory. (B) Sweep angles for the data in (A), relative to the rat’s heading direction (red) or the previous sweep (black). (C) Sweeps in (A) plotted relative to the rat’s location and heading direction. (D) Percentage of sweep pairs that fall on opposite sides relative to the rat’s running direction as a function of the time between sweeps (rat speed > 20 cm/sec; n = 7,285 sweeps). (E) Distribution of sweep angles (absolute value) relative to the rat heading (red) and the previous sweep (black) for the data in (D). (F-G) Distribution of theta periods (F) and sweep distances (G) for the data in (D).

**Figure 2:**
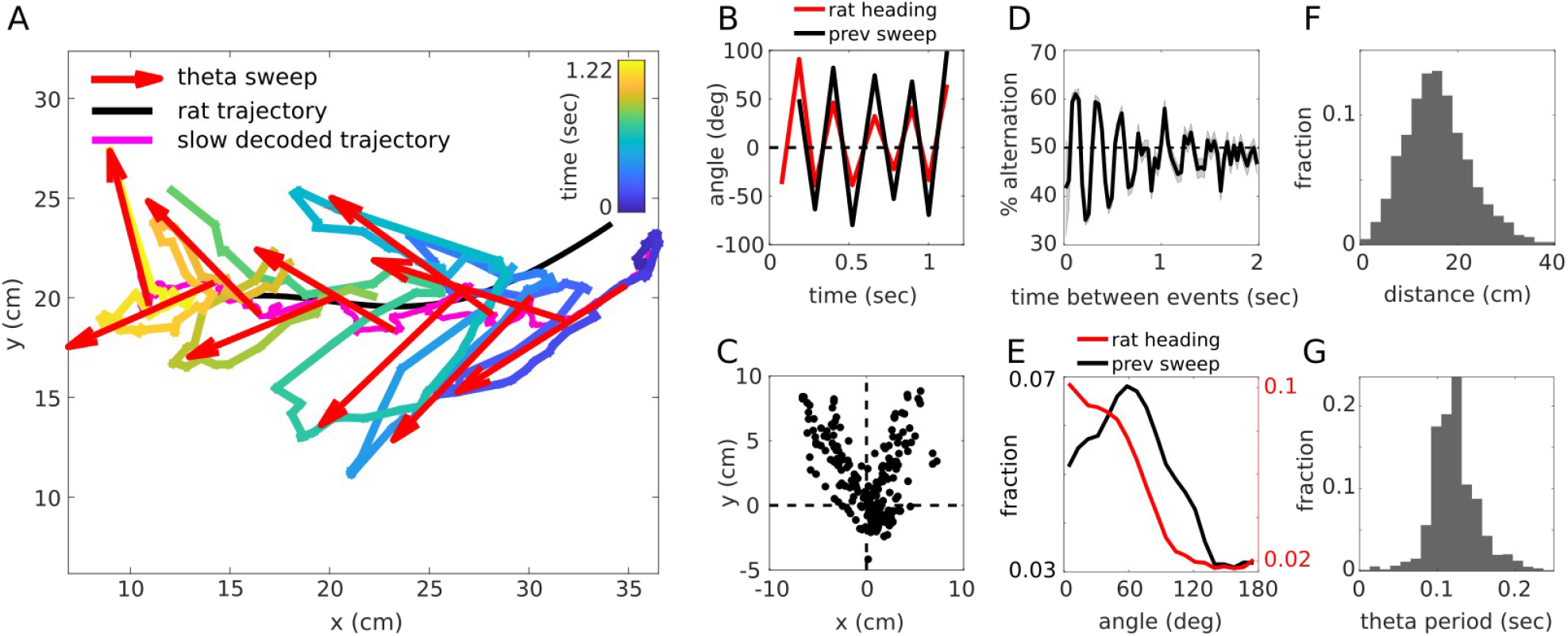
Emergence of 2D alternation in hippocampal data during open field navigation. (A-C) Decoded center-of-mass of hippocampal activity across 10 theta sweeps from rat 1, session 1, depicted as in Figure 1A-C. (D) Percentage of sweep pairs that fall on opposite sides relative to the rat’s running direction as a function of the time between sweeps across one entire recording session (rat speed > 20 cm/sec; n = 2,579 sweeps from rat 1, session 1). (E) Distribution of sweep angles (absolute value) relative to the rat heading (red) and the previous sweep (black) for the data in (D). (F-G) Distribution of theta periods (F) and sweep distances (G) for the data in (D).

Strikingly, the 2D model, which relies on a purely local and thus globally blind control mechanism, elicited sweeps that oriented in the direction of movement while also exhibiting robust alternating structure (**Figure 1A-C**). This alternation obeyed a strict temporal regularity, with the probability of a sweep occurring on the same vs. the opposite side of the rat strongly determined by the time and direction of the previous sweep (**Figure 1D**). The angle of each theta sweep relative to the rat’s heading, and relative to the previous sweep, was precise, peaking around ∼30 deg and ∼60 deg, respectively (**Figure 1E**). Both the temporal regularity and angular precision arose rapidly at the beginning of each bout of movement (**Figure S3**) and were robust across an order-of-magnitude change in network adaptation parameters (**Figure S2**).

We next turned to corroborate our findings using hippocampal data recorded from 2 rats performing a spatial memory task in an open field environment. The task required rats to memorize and repeatedly return a hidden reward location dispensing small amounts of chocolate milk. Theta sequences were computed by applying a Bayesian decoder to the population activity patterns across time (50 msec bins shifted by 5 msec), and then extracting the center-of-mass trajectory of the decoded posterior. The local field potential from a representative channel was used to delineate theta cycles (**Figure S4**). Within each theta cycle, sweep vectors were defined as the line connecting the farthest extent of the decoded center-of-mass trajectory with a slower timescale decoded position (150 msec bins). Compared to the rat’s postion, this slow timescale decoding provided a better reference around which sweeps appeared more regular (**Figure S5**; also pointed out by Vollan et al., 2024). Consistent with previous reports, theta sweeps were readily observed to alternate across the animal’s midline (**Figure 2A-C**). As in the model, this alternation obeyed a strict temporal regularity, with the side of the sweep strongly dependent on the time since the previous sweep (**Figure 2D, Figure S6**). The distribution of angles between successive theta sweeps was strongly peaked around 60 degrees (**Figure 2E, Figure S6**), similar to the model, though the sweep angle relative to the rat’s heading was more irregular and did not show as strong an angular preference.

In sum, we found that our model rapidly and robustly produced theta sweeps that alternated regularly across the animal’s midline, consistent with experimental data reporting theta sweeps in hippocampal-entorhinal data (Vollan et al., 2024) and which we confirmed using a separate hippocampal data set. Our model also provides a biologically plausible, mechanistic account of alternation in 2D theta sweeps, in contrast with more normative approaches based on maximizing information gathering (Ujfalussy & Orban, 2022; Vollan et al., 2024). Notably, our model incorporates circularly symmetric weights and purely local adaptive drive, which means that the model does not inherently possess any direction biases. In other words, sweep alternation and angular precision are emergent features of the model. This also means that sweep alternation need not rely on foot-step driven asymmetries in theta sweeps (Joshi et al., 2023).

Previous work has shown that spike frequency adaptation in a continuous attractor network can successfully explain properties of theta sweeps on linear tracks, including alternation across arms of a T-maze (Chu et al., 2023). Our model supports and extends these findings to 2D, showing that alternating sweeps still emerge robustly and with angular precision in less constrained environments. The adaptation used in our model generalizes in principle to other forms of adaptation like synaptic depression, which, like spike frequency adaptation, acts to destabilize the bump location but through depressing outgoing synapses of active neurons (Tsodyks et al., 1998; York & van Rossum, 1996; Romani & Tsodyks, 2015). Discriminating between these two alternative control mechanisms will be a topic for future work.

Grid cell sweeps have been shown to occur earlier and exhibit more regularity than place cell sweeps (Vollan et al., 2024), suggesting that alternation arises from the MEC and is inherited by the hippocampus. Our model is equivalent to single-bump grid cells models with periodic boundary conditions (Guanella et al., 2007; Fuhs & Touretzky, 2006; Pastoll et al., 2013; Brecht et al., 2014) and is thus capable of explaining grid cell-driven theta sweep alternation in the open field. For grid cell models based on aperiodic, multi-bump networks (Burak & Fiete, 2009; Widloski & Fiete, 2014), adaptation will likely cause different activity bumps to drift independently, leading to the deterioration of the grid pattern. In this case, including secondary excitatory lobes beyond the purely local connectivity (Fuhs & Touretzky, 2006; Widloski & Fiete, 2014) may help stabilize and rigidify the pattern.

Lastly, our experimental finding that theta sweep alternation and angle is also present in hippocampal data while rats performed a goal directed task, in contrast to simple random foraging (Vollan at al., 2024), suggests that sweep alternation regularity is present regardless of task demands.

## METHODS

### I. SIMULATION OF 2D THETA SEQUENCES

#### Model setup

Replays were simulated using a single-bump continuous attractor network (CAN) where the movement of the bump was driven by spike-frequency adaptation (Hansel and Sompolinsky, 1998; Hopfield, 2010; Itskov et al., 2011; Azizi et al., 2013). Given a summed input current *Ii*(*t*) (see below) to the *i*^*th*^ cell, the instantaneous firing rate for that cell was *f*(*I*_*i*_(*t*)), with the neural transfer function *f* given by

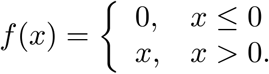

Based on this time-varying input, neurons fired spikes according to a sub-Poisson point process with a coefficient of variance of 0.5 (see Methods section “Sub-Poisson spiking”). The total input to the *i*^th^ cell was a combination of internal and externally-derived inputs (**Figure 1A**), each modulated at theta frequency:

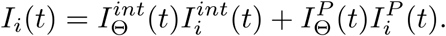

#### Internal inputs

The internal input was given by

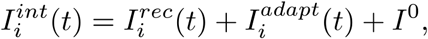

where 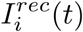 is the recurrent input derived from other cells, 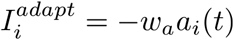 is the adaptive inhibitory input (*w*_*a*_ is the strength of depression) based on adaptation dynamics *a*_*i*_(*t*) (see below), and *I*^0^ is a small positive constant bias common to all cells. The recurrent input was given by

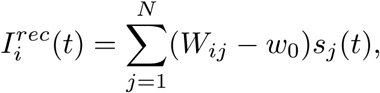

where *W*_*ij*_ are the excitatory recurrent weights from the *j*^th^ to the *i*^th^ cell, *w*_0_ is the strength of recurrent inhibitory feedback, *N* is the number of cells, and *s*_*j*_(*t*) is the *j*^th^ cell’s synaptic activation dynamics (see below). To specify the recurrent weights, cells were organized into a 2D array in the neural sheet, where the location of the *i*^th^ cell was given by **1**_*i*_ For simplicity, the sheet was assumed to be periodically connected, though this was not a necessary component of the model. Let *W*_*ij*_ be a set of translation-invariant symmetric weights with Gaussian shape that depend on the distance between cells in the neural sheet:

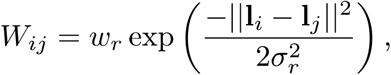

where *w*_*r*_ and σ_*r*_ control the strength and spatial extent of the connectivity, respectively. The synaptic activation dynamics for the *i*^th^ cell, *s*_*i*_(*t*), was given by

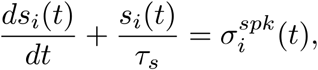

where τ_*s*_ is the synaptic time constant and

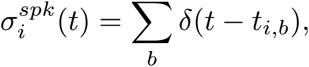

is the cell’s spike train (*t*_*i*,*b*_ specifies the time of the *b*^th^ spike of the cell and the sum is over all spikes of the cell). The adaptation dynamics for the *i*^th^ cell,*a*_*i*_(*t*), was given by

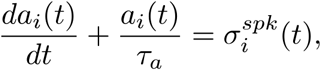

where τ_*a*_ is the time scale of adaptation.

#### Place inputs

The place input into the *i*^th^ cell was given by a 2D Gaussian centered on the rat’s location **x**_*rat*_(*t*) at time *t* with width σ_*rat*_ and amplitude *w*_*rat*_:

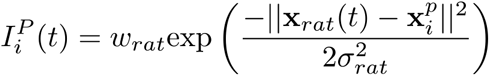

where 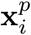 is the cell’s place preference, which were uniformly distributed across a 1 m^2^ square environment. Place preferences were also used to convert coordinates of the bump on the neural sheet to coordinates in rat trajectory space. The rat trajectory was simulated as a correlated random walk as follows. First, a speed profile was specified, which took the form of a sinusoid with period *T*_*run*_ seconds but modified so that the rat dwelled in the rest state (zero velocity) for *T*_*run*_ seconds (**Figure S1B**). Then, for each time step of length Δ*t*, starting from the origin, a spatial displacement was drawn stochastically, with length determined from the speed profile and direction sampled from the normal distribution ℕ( θ_*t* −1,_ σ_θ_) where ℕ is a normal distribution, θ_*t* −1_ is the step direction from the previous time step, and is the σ_θ_ step direction standard deviation (**Figure S1C**). At boundaries, the displacement angle was resampled until the new direction moved away from the wall. As noted, while a periodic network topology was used for the network connectivity, the animal’s trajectory was aperiodic Thus, to avoid the activity bump from unrealistically activating cells on opposite sides of the environment, the “boundaries” used to constrain the rat trajectory were brought in by 10 cm, so that the effective size of the environment was 80 x 80 cm (**Figure S1C**).

#### Theta modulation

Both the internal and place inputs were modulated at theta-frequency (**Figure S1D**), where theta periods were sampled stochastically from the distribution 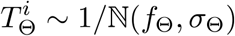, where ℕ is a normal distribution with mean *f*_Θ_ and standard deviation σ_Θ_defined to match empirical observations (compare **Figure 1F** to **Figure 2F**). Note that for simplicity we did not model the speed dependence of the theta period, since all of the analysis was restricted to running periods (speed > 5 cm/sec). The theta modulation 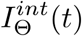 was then defined as follows: For the *i*^th^ theta interval of length 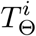 at time 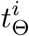, where 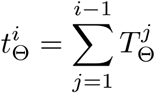 is the cumulative sum of all preceding theta periods,

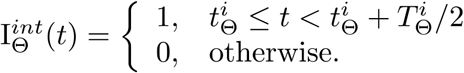

This amounts to a rectified sinusoid with a duty cycle of 50% of the theta period. Likewise, the place inputs were modulated at theta-frequency:

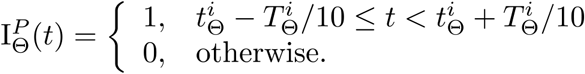

This amounts to a rectified sinusoid with a duty cycle of 20% of the theta period but also shifted in time in order to allow the place inputs to occur briefly around the time of the release of the internally-driven dynamics of the network.

#### Sub-Poisson spiking

To generate spike trains with 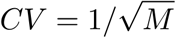, where *CV* is the coefficient-of-variance of the inter-spike interval and *M* is an integer greater than 1, a procedure was borrowed from Burak & Fiete (2009). The interval Δ*t* is first sub-divided into *M* sub-intervals of length Δ*t/M*. Within each of these smaller intervals, Poisson spikes are sampled at a higher mean rate *Mf*(*I*_*i*_(*t*)), after which only the *M*^th^ spike is retained.

#### Theta sequences

Let 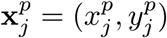 be the coordinates of the *j*^th^cell’s place preference. The center-of-mass of the activity bump in the neural sheet converted to rat trajectory space coordinates was computed as

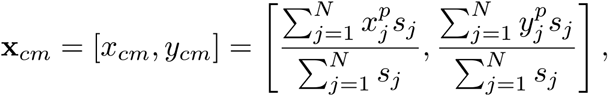

where *s*_*j*_ is the instantaneous synaptic activation of the network. Theta sweeps were extracted as the center-of-mass trajectory during the “on” phase of the internal dynamics (i.e., when 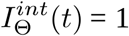). Theta sweep distance was computed as the integrated path length of this trajectory. To compute theta sweep angles, each sweep was characterized by a vector extending from the rat’s position (averaged over the theta cycle) to the position of the sweep at its maximum distance with respect to this point. The angle of the sweep relative to the rat’s heading was defined as the angle subtended by the sweep vector and the rat’s instantaneous velocity vector (averaged over the theta cycle). The angle of the sweep relative to the previous sweep was defined as the angle subtended by the sweep vector and the sweep vector from the previous theta cycle. Sweep alternation was computed as the probability that, for any pair of sweeps, their sweep vectors should point to opposite sides relative to the rat’s heading as a function of the time between sweeps.

#### Simulation parameters

Integration was by the Euler method with step size Δ*t*. Other network parameters were:

| $\Delta t$ | $N$ | $\tau_s$ | $w_r$ | $\sigma_r$ | $w_0$ | $I^0$ | $\tau_a$ | $w_a$ | $w_{rat}$ | $\sigma_{rat}$ | $\sigma_\Theta$ | $T_{run}$ | $T_{rest}$ | $M$ |
| --- | --- | --- | --- | --- | --- | --- | --- | --- | --- | --- | --- | --- | --- | --- |
| 0.0005 | 64 | 0.3 | 60 | 0.1 | 0.07 | 400 | 12 | 4 | 300 | 0.01 | 0.4 | 10 | 10 | 4 |

Temporal parameters (Δ*t*, τ’s and *T*’s) are in units of seconds; amplitudes (*w*’s) are dimensionless; *I*^0^ is in units of spikes/sec; *N* and σ_*r*_ are in units of cells; σ_*rat*_ is in units of meters; σ _Θ_ is in units of Hz.

### II. EXPERIMENTAL DATA ANALYSIS

#### Behavioral paradigm

Neural activity was recorded from dorsal hippocampus (region CA1) of 2 Long-Evans rats (*Rattus norvegicus*; 3-4 months old) performing a goal-directed task in an open field maze (task described in Widloski & Foster, 2022). In brief, rats were trained to search for liquid chocolate available in one of 9 food wells, which alternated on consecutive trials between a learnable fixed location and unpredictable other locations. The identity of the learnable fixed location changed pseudorandomly across sessions. Rats were housed in a humidity and temperature controlled facility with a 12-hour light-dark cycle.

#### Experimental recordings

Rats were implanted with microdrive arrays weighing 40-50 grams and consisting of 64 independent-adjustable tetrodes made of twisted platinum iridium wires (Neuralynx) gold plated to an impedance of 150-300 MOhms. Drive cannulae were implanted bilaterally to target hippocampal dorsal CA1 (−4.13 AP, 2.68 ML relative to bregma) using a surgery protocol described elsewhere (Widloski and Foster, 2022). Tetrodes were slowly lowered to the cell layer over the course of 2-4 weeks, which was identified by the presence (and shape) of strong-amplitude sharp-wave ripples. The rats were allowed 3-4 days recovery, after which behavioral training on the barrier maze task was resumed but without food restriction in their home cages. Food restriction was resumed a week after surgery. All experimental procedures were in accordance with the University of California Berkeley Animal Care and Use Committee and US National Institutes of Health guidelines.

#### Place fields

Spikes were extracted from channel LFP’s (sampled at 30 kHz) using Spike Gadgets Trodes software and clustered automatically using Mountainsort (Chung et al., 2017) and merged across sessions using the *msdrift* package. Additional cluster mergings across sessions was performed manually based on similarity of waveform. Clusters were accepted if noise overlap < 0.03, isolation > 0.95, peak SNR > 1.5 (Chung et al., 2017) and had passed a visual inspection. Place fields were determined for each cluster as follows. First, rat position was tracked using automated software from Spike Gadgets and sampled at 30 Hz. Position and velocity were smoothed using a Butterworth filter (second order with a cutoff frequency of 0.1 samples/s using the butter function in Matlab, selected to give reasonable smoothing to the rat’s trajectory). For each spike that occurred during run (rat speed > 5 cm/sec), the rat position was found through linear interpolation (*interp1* in Matlab). The unsmoothed rate map for the *i*^th^ cell was defined as

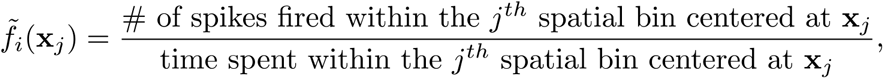

where positions were binned with 2 cm square bins. Smoothed rate maps, denoted as *f*_*i*_ (x_*j*_), were computed by first setting unvisited bins to zero and convolving the rate maps with a 2D isotropic Gaussian kernel (8 cm standard deviation (SD)). Spatial information (bits/spike) for the *i*^th^ cell was defined as

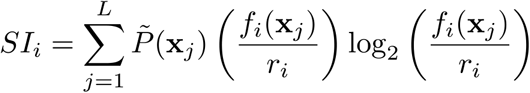

where L is the number of spatial bins,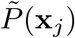 is the probability of the rat or replay being at the *j*^th^ spatial bin, and 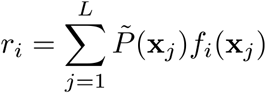 is the cell’s mean firing rate. *Place cells* were identified as having place fields with *r*> 0.01 Hz and *SI*> 0.5 bits/sec.

#### Bayesian decoding

Let *k*_*i*_ be the number of spikes emitted by *i*^th^ the place cell in a given time bin of duration τ. The posterior probability at bin x_*j*_ conditioned on the activity vector 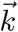 (with the *i*^th^ element as *k*_*i*_) is given by Bayes rule (assuming Poisson spiking noise statistics, independence between cells, and a uniform spatial prior):

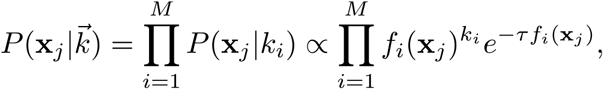

where *M* is the number of neurons. A uniform prior was used for the purposes of making minimal assumptions about the location of the decoded positions. The posterior probability was computed for all bin locations x_*j*_ where 1≤ *j* ≤ *L* and *L* is the total number of spatial bins. Define 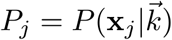 and let x_*j*_=(*x*_*j*,_*y*_*j*_) be the components of the *j*^th^ spatial bin. The components of the *posterior center-of-mass* (COM) were given by

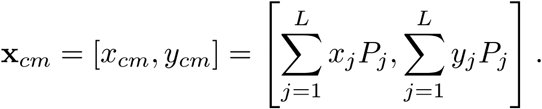

The size of the time bins, τ, used to decoded theta sweeps was 50 msec, shifted by 5 msec increments. A slower time-scale decoding using 150 msec time bins, also shifted by 5 msec, was also used to compute a reference point to measure the sweeps against (see below).

#### Theta sequences

For each session, the LFP from a single representative tetrode channel exhibiting strong theta power was bandpassed in the theta range (4-12 Hz), then Hilbert transformed in order to determine theta phase as a function of time. Since theta phase can vary depending on the position of the tetrode within the hippocampus, theta phase was realigned such that the distance between the decoded center-of-mass and the rat position was maximal at zero phase (**Figure S4**). Theta sweeps were extracted as the decoded center-of-mass trajectory within the theta phases of 180 and 360 degrees, which corresponds to when the sweep is, on average, closest to and farthest from the rat’s position, respectively. Theta sweep distance was computed as the integrated path length of this trajectory. To compute theta sweep angles, each sweep was characterized by a vector extending from the slow time-scale decoded location (averaged over the theta cycle) to the position of the sweep at its maximum distance with respect to this point. The angle of the sweep relative to the rat’s heading was defined as the angle subtended by the sweep vector and the rat’s instantaneous velocity vector (averaged over the theta cycle). The angle of the sweep relative to the previous sweep was defined as the angle subtended by the sweep vector and the sweep vector from the previous theta cycle. Sweep alternation was computed as the probability that, for any pair of sweeps, their sweep vectors should point to opposite sides relative to the rat’s heading as a function of the time between sweeps.

## ACKNOWLEDGEMENTS

The authors would like to thank Sandro Romani for an insightful conversation during a long plane ride across the Atlantic ocean.

## SUPPLEMENTAL INFORMATION

**Figure S1:**
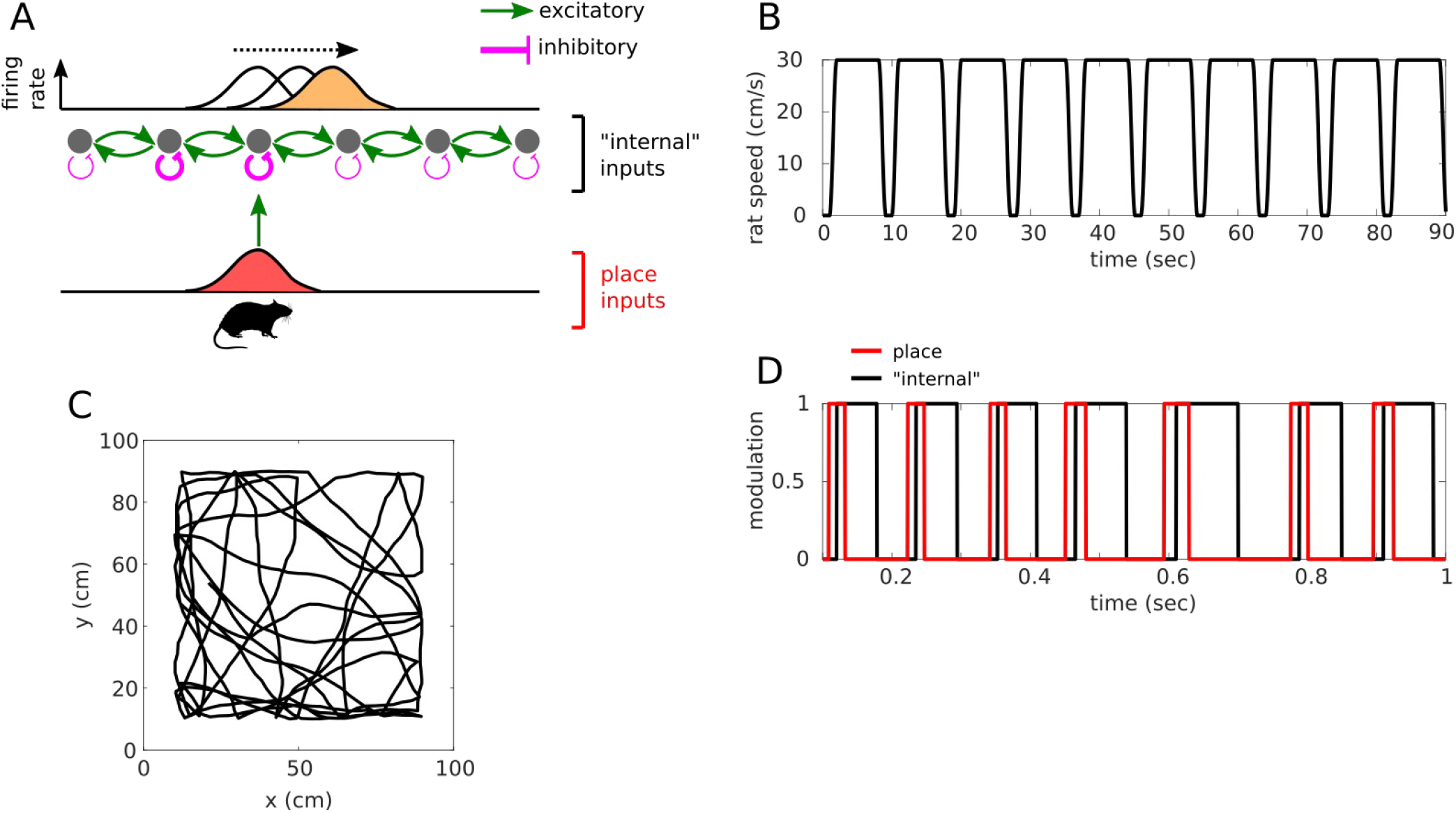
Model setup. (A) Schematic showing model components. Top: Network connectivity is composed of local excitation (green arrows) and global inhibition (not shown), which supports the formation of activity bump (yellow). Adaptive inhibitory feedback (pink lines) based on each cell’s instantaneous firing rate destabilizes the bump and causes it to move. Bottom: The place inputs derived from the rats trajectory (bottom) reset the bump to the rat’s location at the beginning of each theta cycle. (B-C) Rat speed profile (B) and simulated rat trajectory (C) across 6 movement bouts. (D) Theta modulation of “recurrent” and place inputs across 7 theta cycles (see Methods).

**Figure S2:**
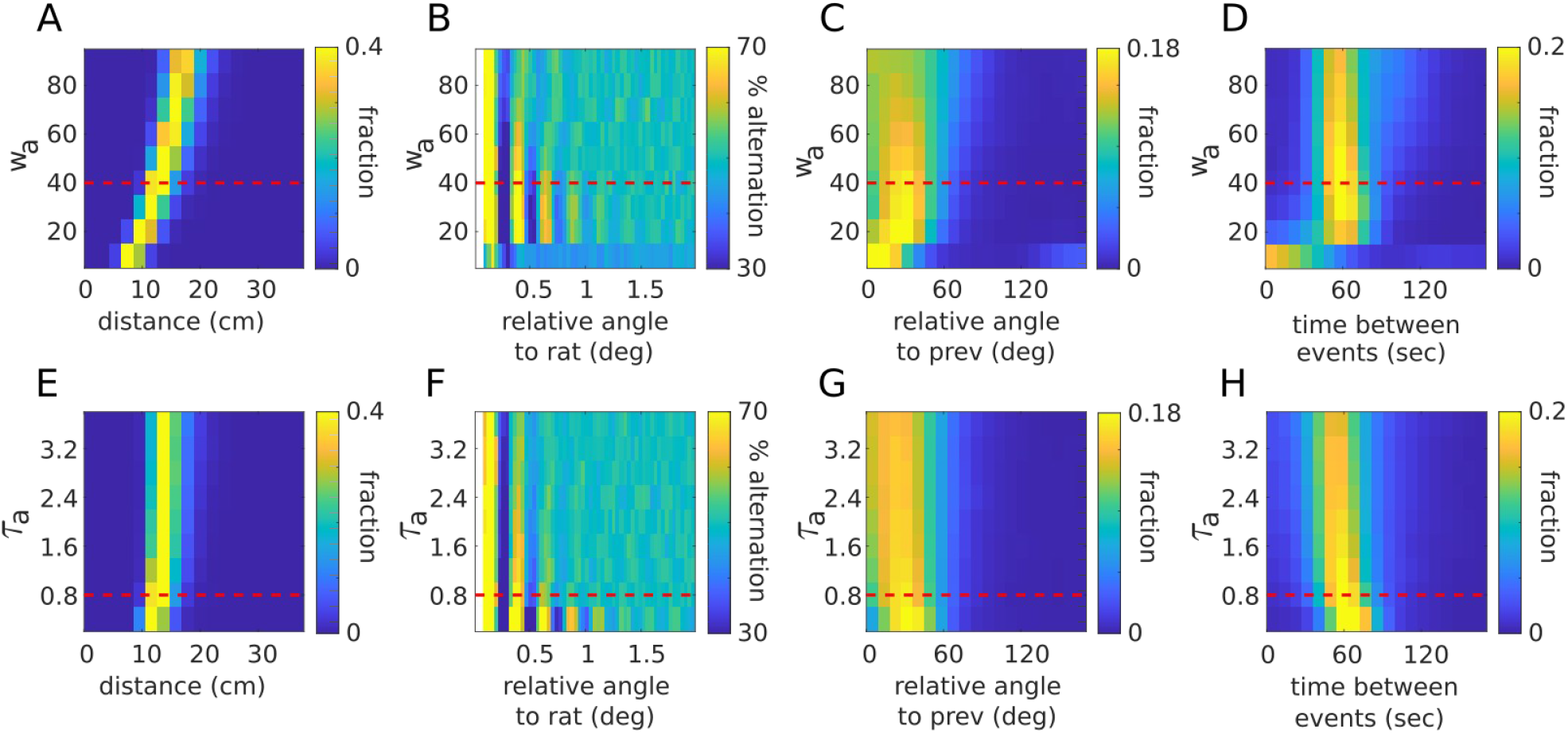
Robustness of temporal regularity and angular precision to network parameters. (A) Distribution of sweep distances (rows) as a function of the strength of adaptation,w_a_, along the y-axis while keeping other parameters fixed. Dashed red line indicates the value of w_a_ used for Figure 1. (B-D) Same as in (A) except plotting the percentage of sweep pairs that fall on opposite sides relative to the rat’s running direction as a function of the time between sweeps (B; as in Figure 1D), distribution of sweep angles relative to the rat heading (C, as in Figure 1E, red), and the distribution of sweep angles relative to the previous sweep (D, as in Figure 1E, black). (E-F) Same as (A-D) except varying the time constant of adaptation, τ_a_, along the y-axis while keeping other parameters fixed.

**Figure S3:**
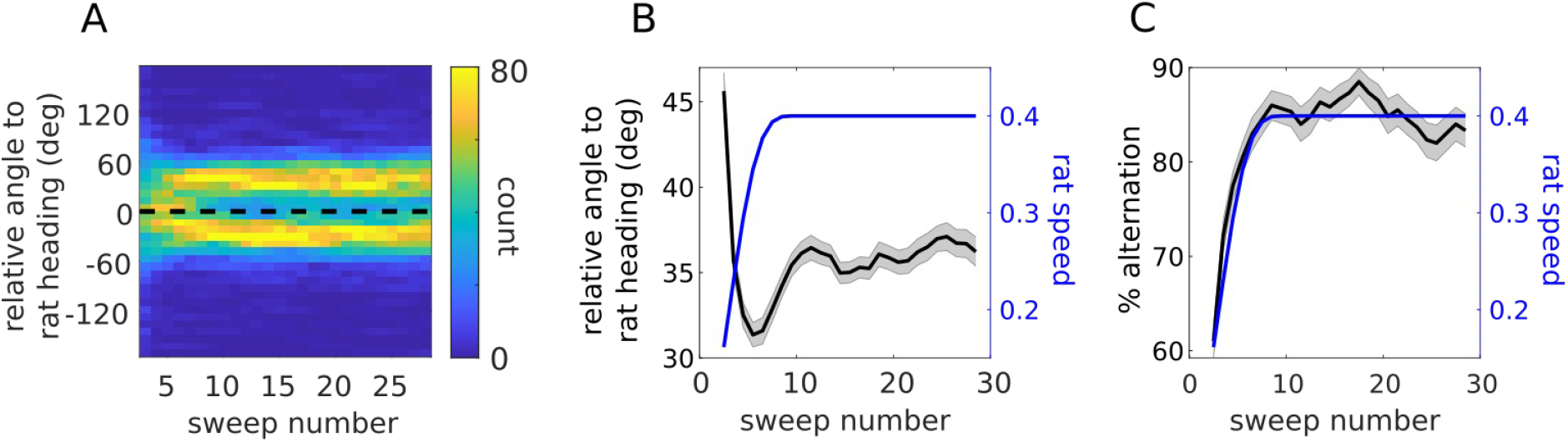
Sequence alternation emerges rapidly at the beginning of each movement bout. (A) Distribution of sweep angles (columns) relative to the rat’s heading as a function of sweep number along the x-axis, starting from the beginning of each movement bout. (B) Average sweep angle relative to the rat’s heading (black) and average rat speed (blue) as a function of sweep number within a movement bout. (C) Average number of sweeps that are preceded by and follow a sweep in the opposite direction (i.e., exhibit perfect alternation), as a function of sweep number within a movement bout (n = 200 bouts; traces show mean ± sem)

**Figure S4:**
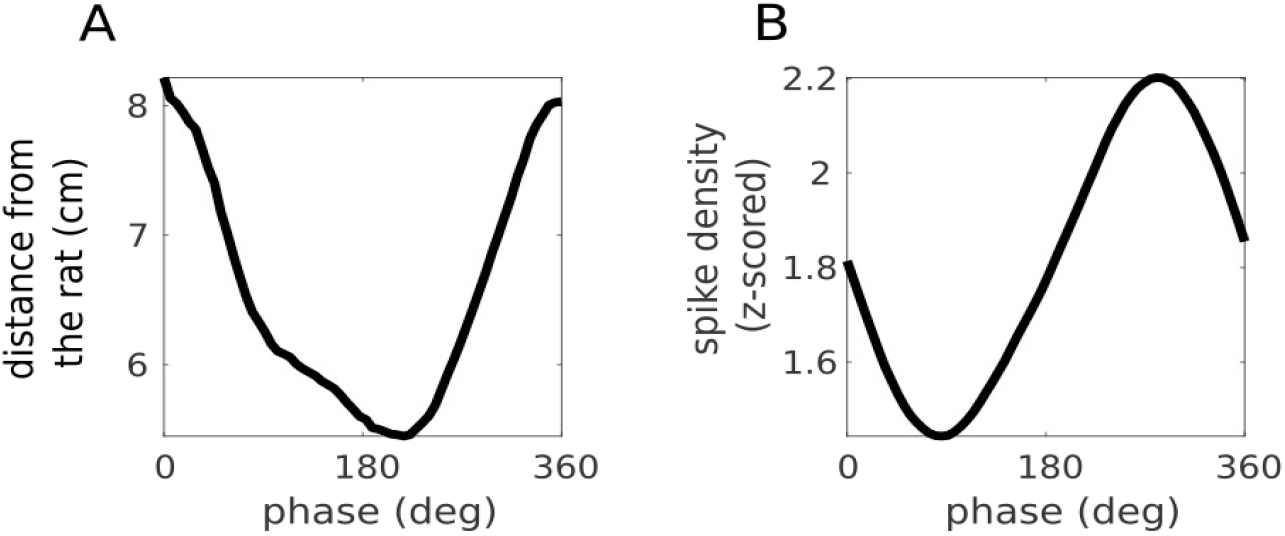
Sweep distance and spike density as a function of theta phase. (A) Distance between the center-of-mass of the decoded posterior and the rat’s location as a function theta phase, averaged across all theta cycles from rat 1, session 1 (rat speed > 5 cm/sec). Distance was first interpolated across equi-spaced phases of theta before averaging. Theta phase has been phase shifted so that the curve peaks at zero phase. (B) Spike density as a function of theta phase. Spike density was computed by first summing the total number of spikes from all clusters in 1 msec non-overlapping time bins, then smoothed through convolution with a Gaussian kernel (80 msec) and z-scored. Spike density as a function of theta phase was computed by interpolating and averaging, as in (A).

**Figure S5:**
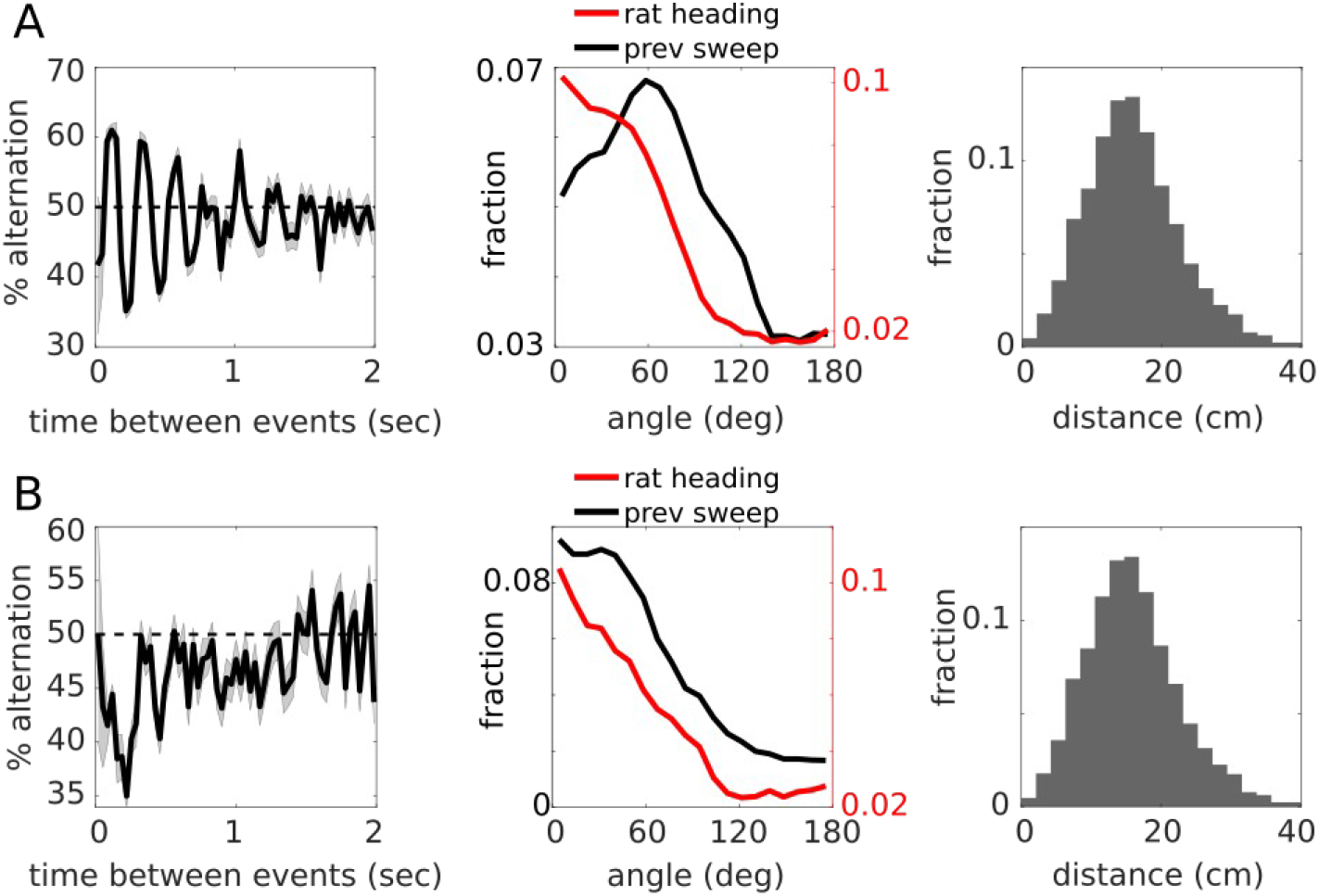
Theta sweeps measured with respect to different reference frames. (A-B) Depicted the same as in Figure 2D-E & G, where theta sweep angles have been measured relative to the slow time-scale decoding (A, copied from Figure 2D-E & G), or the rat’s position (B).

**Figure S6.**
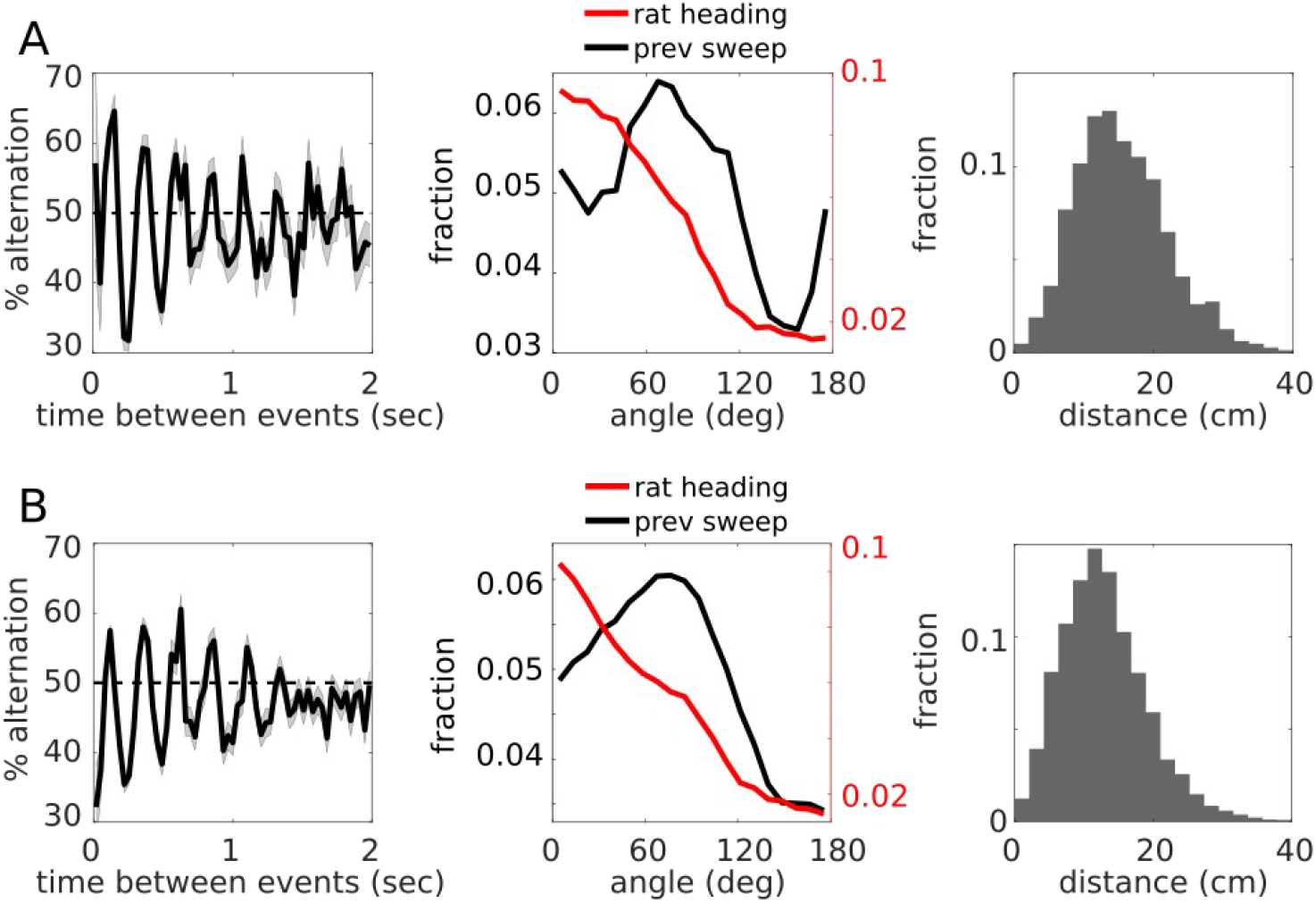
Main experimental findings are recapitulated across sessions and across rats. (A-B) Depicted the same as Figure 2D-E & G, except for rat 1, session 2 (A), and rat 2, session 1 (B).

